# Maintaining trunk neural crest cells as crestospheres

**DOI:** 10.1101/391599

**Authors:** Sofie Mohlin, Ezgi Kunttas, Camilla U. Persson, Reem Abdel-Haq, Aldo Castillo, Christina Murko, Marianne E. Bronner, Laura Kerosuo

## Abstract

Neural crest cells have broad migratory and differentiative ability that differs according to their axial level of origin. However, their transient nature has limited understanding of their stem cell and self-renewal properties. While an *in vitro* culture method has made it possible to maintain cranial neural crest cells as self-renewing multipotent crestospheres (Kerosuo et al., 2015), these same conditions failed to preserve trunk neural crest in a stem-like state. Here we optimize culture conditions for maintenance of trunk crestospheres, comprised of both neural crest stem and progenitor cells. Trunk crestospheres display elevated expression of neural crest cell markers as compared to those characteristic of neural tube or mesodermal fates. Moreover, trunk crestospheres have increased expression of trunk-related markers as compared to cranial genes. Finally, we use lentiviral transduction as a tool to manipulate gene expression in trunk crestospheres. Taken together, this method enables long-term *in vitro* maintenance and manipulation of trunk neural crest cells in a premigratory stem or early progenitor state to probe the mechanisms underlying their stemness and lineage decisions.

**Highlights:**

- Trunk-derived multipotent neural crest stem cells can be cultured as crestospheres
- Trunk-derived crestospheres require different conditions than cranial
- Trunk crestospheres consist of neural crest stem and progenitor cells
- Trunk crestospheres can be efficiently transduced using lentiviral vectors

## Introduction

The neural crest is a multipotent stem cell population that forms essential structures of the vertebrate body. Arising within the neuroectoderm after gastrulation, premigratory neural crest cells within the dorsal neural tube express numerous transcription factors including *FOXD3, TFAP2* and *SOXE* (Khudyakov and Bronner-Fraser, 2009). Around the time of neural tube closure, neural crest cells delaminate from the neural tube by undergoing an epithelial-to-mesenchymal transition (EMT), a feature shared with metastatic cancer cells, and then migrate extensively to populate distant sites in the embryo. In their final sites, neural crest cells differentiate into more than thirty different cell types, the range of which varies according to their anterior to posterior axial level of origin.

Elegant quail-chick grafting experiments (Ayer-Le Lievre and Le Douarin, 1982) have shown that different populations of neural crest cells arise at different levels of the body axis designated from anterior to posterior as cranial, vagal, trunk and sacral. These four subdivisions all share the ability to form cells of the peripheral nervous system and melanocytes, but also give rise to axial level-specific features. For example, facial bone/cartilage arise from cranial neural crest (Ayer-Le Lievre and Le Douarin, 1982) whereas chromaffin cells of the adrenal medulla only arise from the trunk (Vega-Lopez et al., 2018).

The stem cell properties of neural crest cells make them a cell type of interest in regenerative medicine as well as a possible cell of origin for neural crest-derived tumors and birth defects (Vega-Lopez et al., 2018), highlighting the importance of understanding the regulatory mechanisms underlying neural crest stemness. To address this, we recently reported establishment of crestospheres for cranial neural crest cells. These are self-renewing multipotent primary neural crest cultures derived from either chick embryos or human embryonic stem cells (Kerosuo et al., 2015). Crestospheres maintain neural crest cells as premigratory neuroepithelial cells and differ from previous culture techniques which induce spontaneous differentiation (Baroffio et al., 1988; Curchoe et al., 2010; Lee et al., 2007; Stemple and Anderson, 1992; Trentin et al., 2004). Under crestosphere conditions, premigratory neural crest cells retain multipotency on a clonal level (Kerosuo et al., 2015). Given that there are known axial level differences between neural crest populations and different signaling cascades that determine the anterior to posterior patterning of the vertebrate body, it is not surprising that the established cranial crestosphere conditions do not support maintenance of trunk-derived neural crest cells.

To circumvent this problem, here, we provide conditions that enable long term growth of trunk crestospheres as epithelial spheres under stem cell-promoting culture conditions. Trunk crestospheres express general markers of the premigratory neural crest as well as genes specific to the trunk level, as shown by quantitative PCR, *in situ* hybridization and immunohistochemistry. Importantly, trunk neural crest-derived crestospheres can be efficiently transduced with fluorescently labelled lentiviral vectors. Thus, they hold great promise as an *in vitro* model for examining trunk neural crest-associated birth defects and malignancies such as the childhood tumor form neuroblastoma.

## Methods

### Chick embryos

Chick embryos were obtained from commercially purchased fertilized eggs and incubated at 37.5°C/100°F until the desired developmental stages. For trunk cultures, we used stage 13-/14+ (17-21 somite stage) embryos. For the cranial cultures used as controls, stage 8-/9 (4-7 somite stage) embryos were used, staged according to the criteria of Hamburger Hamilton (HH) (Hamburger and Hamilton, 1951). In a few cases, we tested cranial cultures from later stages when most of the neural crest has already migrated from the neural tube (i.e. HH10+).

### Neural tube dissection

Embryos at designated somite stages were collected from the eggs by using Whatman filter paper with hole-punches. Embryos are placed in the center and transferred to Ringer’s balanced salt solution (Solution-1: 144g NaCl, 4.5g CaCl•2H_2_O, 7.4g KCl, ddH_2_O to 500ml; Solution-2: 4.35g Na_2_HPO_4_•7H2O, 0.4g KH_P_O_4_, ddH_2_O to 500ml (adjust final pH to 7.4). Embryos were detached from the filter paper and placed ventral side up to cut out the endoderm by “cutting from the ventral midline”. Neural tubes from respective axial levels were carefully dissected out, and all neighboring mesoderm and notochord tissue was removed. Isolated neural tubes were transferred to a small volume (50 μl) of sterile PBS. For cranial-derived cultures used as controls, the very anterior tip was excluded, and the neural tube was dissected until the first somite level as previously described (Kerosuo et al., 2015). For trunk-derived cultures, the neural tube was dissected between somites 10-15, and pools of neural tubes from 4 - 6 embryos were used for each culture.

### Cell culture

Pooled neural tubes were mechanically dissociated (~ 30 times until clumps of ~ 50-100 cells were formed) and transferred to NC medium (DMEM with 4.5g/L glucose, 7.5% chick embryo extract, 1X B27, basic fibroblast growth factor (bFGF, 20 ng/ml), insulin growth factor -I (IGF-I, 20 ng/ml), retinoic acid (RA, 60nM for cranial and 180nM for trunk, respectively), and with or without 25 ng/ml BMP-4 in low-adherence T25 tissue culture flasks (2 ml (cranial) or 1.5ml (trunk) per flask in upright position). See Table 1 for complete culture conditions. During the course of 7-14 days, the total volume of media was incrementally increased to a maximum of V_tot_=10ml. Due to rapid degradation, retinoic acid and BMP-4 were re-added to the culture medium volume every 2-3 days, and the spheres were also gently dispersed by pipetting up and down ~10-20 times against the culture flask wall every 2-3 days.

**Table 1.** Culture medium optimized for trunk crestospheres.

Crestospheres - Trunk
| Medium (NC) | C<sub>Final</sub> |
| --- | --- |
| DMEM (with P/S) |  |
| Chicken embryo extract | 7.5% |
| B27 | 1X |
| IGF-I | 20 ng/ml |
| FGF | 20 ng/ml |
| Retinoic Acid | 180 nM |
| BMP-4 | 25 ng/ml |
| DMEM | Corning #10-013-CVR, with 4.5g/L Glucose |
| Chick Embryo Extract (CEE) | MP Biomedicals #092850145 |
| B27 | Life Technologies #12587-010 – 50X |
| Human recombinant IGF-I | Sigma Aldrich #SRP3069 – in ddH<sub>2</sub>O+5% Trehalose in PBS (1:1) |
| Human recombinant bFGF | Peprtech – in Tris 5mM pH7.6 |
| Retinoic acid | Sigma Aldrich #R2625 - 10 mM in DMSO |
| BMP-4 | Peprtech #120-05 – in citric acid (pH3.0):PBS with 2mg/ml BSA (1:1) |

### RNA extraction and quantitative real-time PCR

Total RNA was extracted using the RNAqeuous Micro Kit (Ambion) and eluted in 20μl elution solution. cDNA synthesis using random primers and qRT-PCR was performed as previously described (Mohlin et al., 2015). Relative mRNA levels were normalized to expression of two reference genes (*18S, 28S*) using the comparative Ct method (Vandesompele et al., 2002). See Table 2 for primer sequences.

**Table 2.**
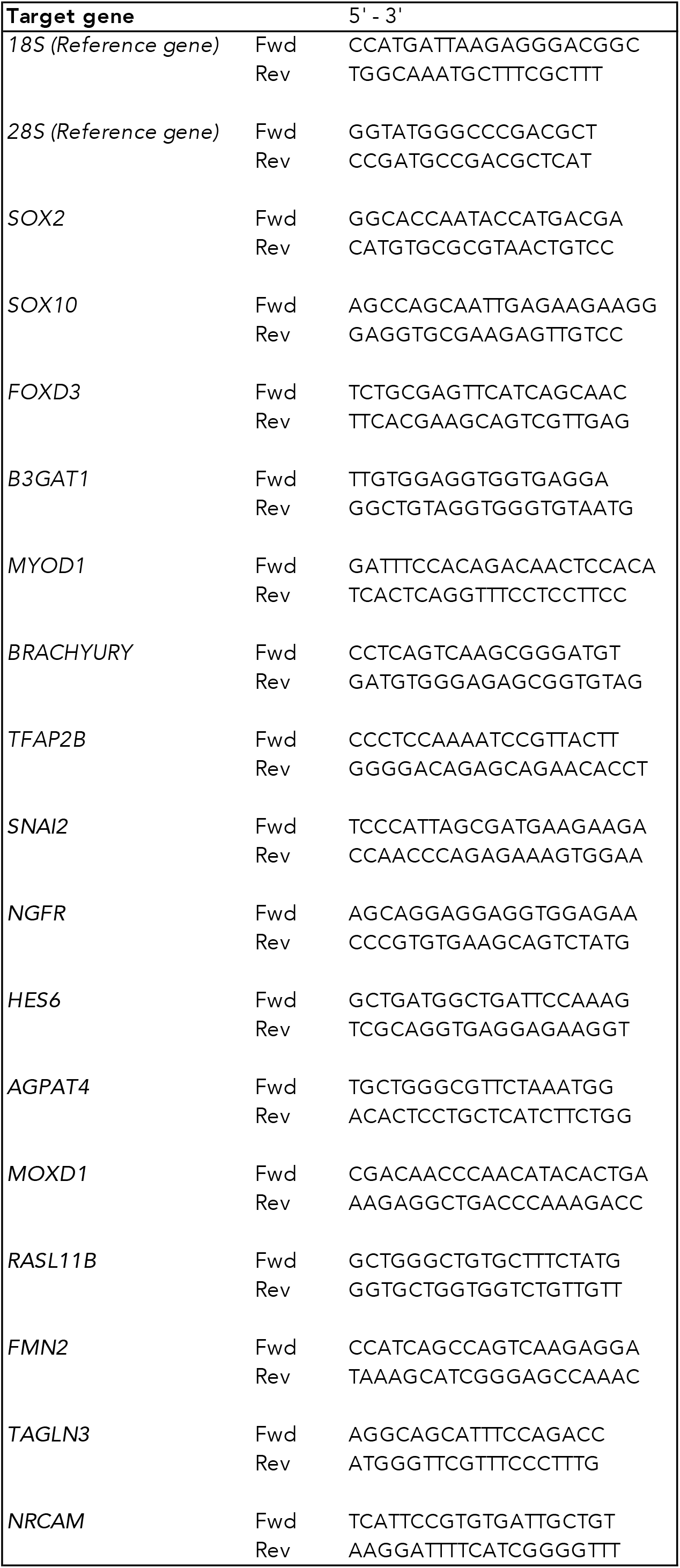
Primer sequences used for quantitative PCR.

### Whole mount in situ hybridization of embryos and crestospheres

For whole mount *in situ* hybridization, embryos were fixed overnight in 4% paraformaldehyde (PFA) at +4°C, washed in PBS containing 0.1% Tween 20 with 0.1% DEPC (DEPC-PBT), dehydrated in a methanol (MeOH)/PBT series at room temperature and kept in 100% MeOH at −20°C until use (days or even months). *In situ* hybridization was performed as previously described (Acloque et al., 2008); briefly, embryos were rehydrated back to 100% PBT and prehybridized in hybridization buffer for 2 hours at 70°C. Embryos were then hybridized with Digoxigenin (DIG)-labeled probes overnight at 70°C. The next day, embryos were washed in prehybridization buffer multiple times before switching to Maleic Acid Buffer (MAB) with 0.1% Tween 20 (MABT). Embryos were then blocked in MABT solution with 10% Boehringer Blocking Reagent and 10% Sheep Serum for 2 hours and incubated with an anti-DIG antibody (1:2000) (Roche) in blocking solution overnight at 4°C. On day 3, embryos were washed in MABT throughout the day and then switched into Alkaline phosphatase buffer (NTMT; 100mM NaCl, 100mM Tris-Cl (pH 9.5), 50mM MgCl2, 1%Tween-20) before visualizing the signal using nitroblue tetrazolium (NBT) and 5-Bromo-4-chloro-3-indolyl phosphate (BCIP) solutions (Roche). Embryos were fixed in 4% PFA for 20 minutes when they reached the desired state and dehydrated in MeOH to be stored at −20°C. Images of whole mount embryos were taken using an Axioskop2 (Zeiss) microscope equipped with Axiovision software, and a region between somites 10-15 was dissected and embedded in blocks of gelatin for transverse sectioning at 20 μm using a cryostat.

Similarly, crestospheres were fixed in 4% PFA for 30 minutes at RT or overnight at +4°C and washed in DEPC-PBT. Samples were then gradually dehydrated by bringing them to 100% MeOH and kept at −20°C for a minimum of 2 hours (or long-term for months). *In situ* hybridization was performed as described above (Acloque et al., 2008). Avian DIG-labelled hybridization probes for *SOX10, FOXD3, PAX7, SNAI2, SOX2* and *SOX9* were cloned by using chicken cDNA as previously described (Khudyakov and Bronner-Fraser, 2009).

### Whole mount immunohistochemistry

Crestospheres were fixed in 4% paraformaldehyde for 15 minutes at RT, washed three times in TBS-T (TBS + Ca^2+^ supplemented with 0.1% Triton-X) and blocked in 10% goat serum in TBS-T for 4 hours at RT. Crestospheres were incubated with primary antibody (PAX7, Hybridoma Bank, 1:10) diluted in block solution over night at +4°C. After two times washing for 30 minutes in TBS-T, crestospheres were incubated with secondary antibody (goat antimouse Alexa Fluor-594 [A11032]; 1:1000) and DAPI (Dako [D3571], 1:3000 diluted in blocking solution) for 4 hours at RT. Crestospheres were washed twice in TBS-T for 30 minutes before imaged mounting the crestospheres in cavity slides using the Slowfade anti-fade Kit (Dako, S2828).

### Lentiviral transduction

Crestospheres were dissociated into single cells / small clusters by using Accutase (Sigma Aldrich; incubation at 37°C for 30 minutes) and seeded at high density in 150 μL culture medium in 24-well plates. Cells were directly transduced with increasing concentrations of pCIG3-GFP (pCMV-IRES-GFP version 3, kind gift from Felicia Goodrum (Addgene plasmid # 78264) (Caviness et al., 2014)). Medium was changed twice a day for four consecutive days. GFP positivity was analyzed 96 hours post-transduction using an Olympus inverted fluorescence microscope. Virus titers were optimized and calculated using sphere growing neuroblastoma patient-derived xenograft cells (Persson et al., 2017) due to the difficulty in obtaining pure single cell suspensions of trunk crestosphere cultures. Two titers were tested; 2.1*10^8 TU/ml and 2.3*10^8 TU/ml; TU, transducing units.

## Results & discussion

### Optimization of the correct trunk premigratory axial level

Cranial crestospheres grow as three-dimensional spheres under conditions optimized for neural tubes from chick embryos as well as human embryonic stem (ES) cells; these spheres retain their self-renewal capacity and multipotency for up to seven weeks in culture (Kerosuo et al., 2015). Here, we sought to devise modifications that would enable growth and maintenance of premigratory neural crest cells from more posterior axial levels to provide a tool for studies on trunk neural crest stemness as well as trunk-derived neurocristopathies.

To ensure that the proper stage from which to derive crestosphere cultures was used, we performed *in situ* hybridization on whole mount embryos using neural crest markers *SOX10, FOXD3, PAX7, SOX9, SNAI2* and the neural marker *SOX2* (Figure 1A-F). As expected the endogenous expression patterns showed that the trunk neural crest was premigratory at the level adjacent to somites 10-16 in stage HH13-/14 embryos.

**Figure 1.**
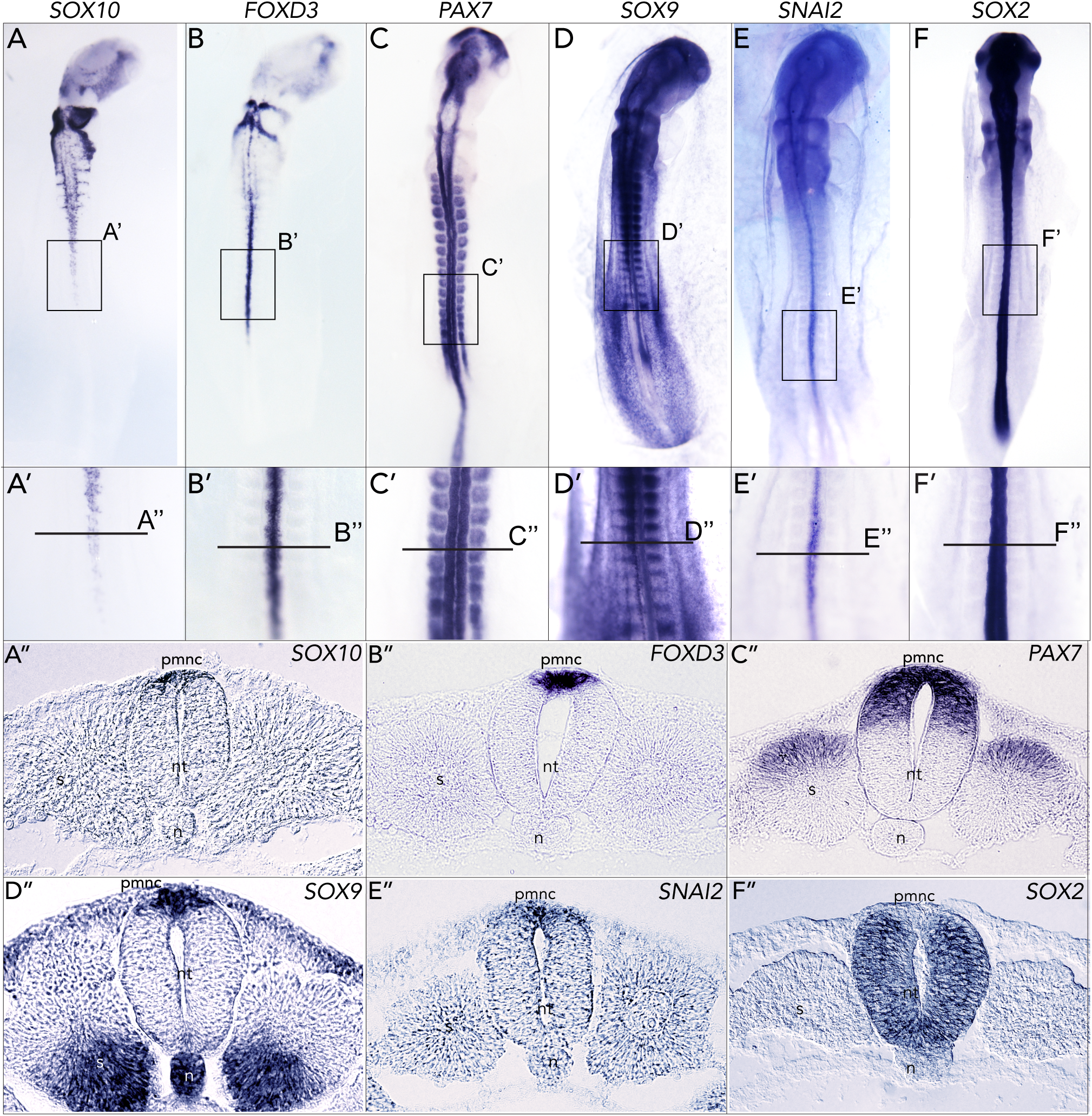
Endogenous expression of premigratory neural crest markers in the posterior axial levels. **(A-F)** Whole mount *in situ* hybridization of HH13-14 embryos. A’-F’ Inserts showing the neural tube at the respective somite level that was dissected for trunk crestosphere cultures. (A”-F”) Transverse sections of the neural tubes at the level that was used for trunk crestosphere cultures.

We tested our cranial crestosphere protocol on dissociated cell clusters of trunk neural tubes derived from chick embryos at several stages (HH10+, HH12, HH13/14), and confirmed that 17-21 somite stage (HH13/14) embryos was the optimal developmental stage for trunk crestosphere culture establishment (Figure 2A). With HH13/14 embryos, spheres formed rapidly after tissue dissociation and expanded accordingly during *in vitro* culture, whereas trunk crestospheres derived from HH10+ or HH12 embryos grew slower and expressed lower levels of neural crest markers (data not shown).

**Figure 2.**
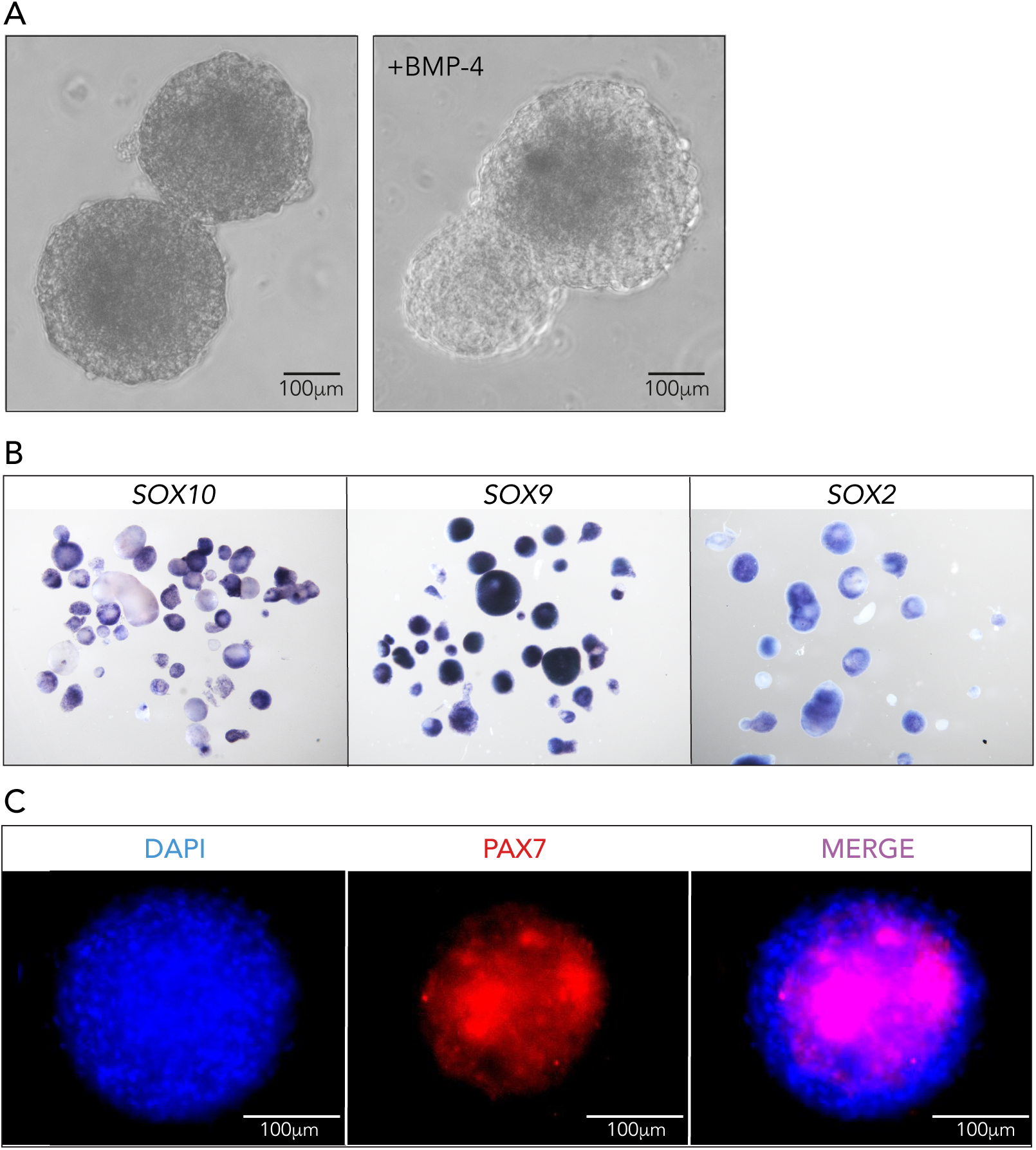
Trunk crestospheres are enriched for neural crest marker expression. **(A)** Trunk crestospheres cultured without or with BMP-4 *in vitro* grow in similar fashion. **(B)** *In situ* hybridization for *SOX10* and *SOX9*. Trunk crestospheres also show weak expression of the neural stem cell marker *SOX2*. **(C)** PAX7 immunopositive neural crest cells in a trunk crestosphere.

### Trunk neural crest cells do not survive in conditions optimized for cranial crestospheres

Although it was initially possible to establish trunk crestospheres under culture conditions optimized for cranial crestospheres (Kerosuo et al., 2015), trunk crestospheres failed to thrive and expand long term. All cultures resulted in less than 25 spheres each during the entire 3 week culture period (n=4). In comparison, all three simultaneously established cranial cultures produced 100+ spheres (n=3).

Given the known importance of retinoic acid (RA) in posteriorizing the body axis as well as in caudal neural crest development (Kudoh et al., 2002; Martinez-Morales et al., 2011; Retnoaji et al., 2014), we increased the amount of RA in the culture medium from 60nM (Kerosuo et al., 2015) to 180 nM. Visual observation of sphere growth suggested that 180nM of RA was optimal for successful expansion of trunk-derived crestospheres. All cultures resulted in more than 50 to 100 spheres (n=6). Despite the additional supply of RA, trunk-derived crestosphere cultures continued to grow more slowly than cranial-derived cultures. We therefore optimized the starting culture volume by further decreasing it from 2 ml used for cranial crestospheres (Kerosuo et al., 2015) to 1.5 ml NC medium. This decrease in starting culture volume had a positive impact on trunk crestosphere expansion, suggesting that the crestospheres condition their own medium by producing additional essential growth factors. The number of spheres in each culture reached ~50 in less than 48 hours (n=3), equivalent to that observed for cranial-derived spheres under optimal cranial crestosphere conditions. In similarity to previously established cranial crestospheres (Kerosuo et al., 2015), trunk crestosphere cultures grew and expanded *in vitro* for 6-7 weeks, after which they were still viable but virtually stopped to proliferate.

### Trunk crestospheres are enriched for neural crest cells

Next, we performed *in situ* hybridization to test expression of neural crest markers in our established sphere cultures. Trunk crestospheres showed strong expression of the neural crest markers *SOX10* and *SOX9* (Figure 2B), and as expected, we also detected some weaker level expression of the neural stem cell marker *SOX2* (Figure 2B). Trunk crestospheres also contained PAX7 immunopositive cells, further confirming the enrichment of neural crest cells (Figure 2C).

### Trunk crestospheres are not enriched for mesodermal fates

To further characterize gene expression patterns of the trunk crestosphere cultures, and to ensure lack of contamination from neighboring mesodermal cells, we used quantitative PCR (qPCR) to address expression levels of neural crest markers relative to markers that reflect the presence of mesodermal or mesenchymal fates. Consistent with *in situ* hybridization results, trunk crestospheres display increased expression of neural crest markers *SOX10* and *FOXD3*, as well as *B3GAT1* (encoding the enzyme required for formation of the HNK1-epitope; expressed in chicken neural crest), as compared to expression levels in whole embryos. Meanwhile, the relative expression levels of mesodermal/mesenchymal genes *MYOD1* and *T* (T <u>brachyury</u> transcription factor) were low (Figure 3A). These results confirm the primarily neural crest identity of trunk crestospheres and rule out contamination by mesoderm- and/or mesenchymal-derived cells.

**Figure 3.**
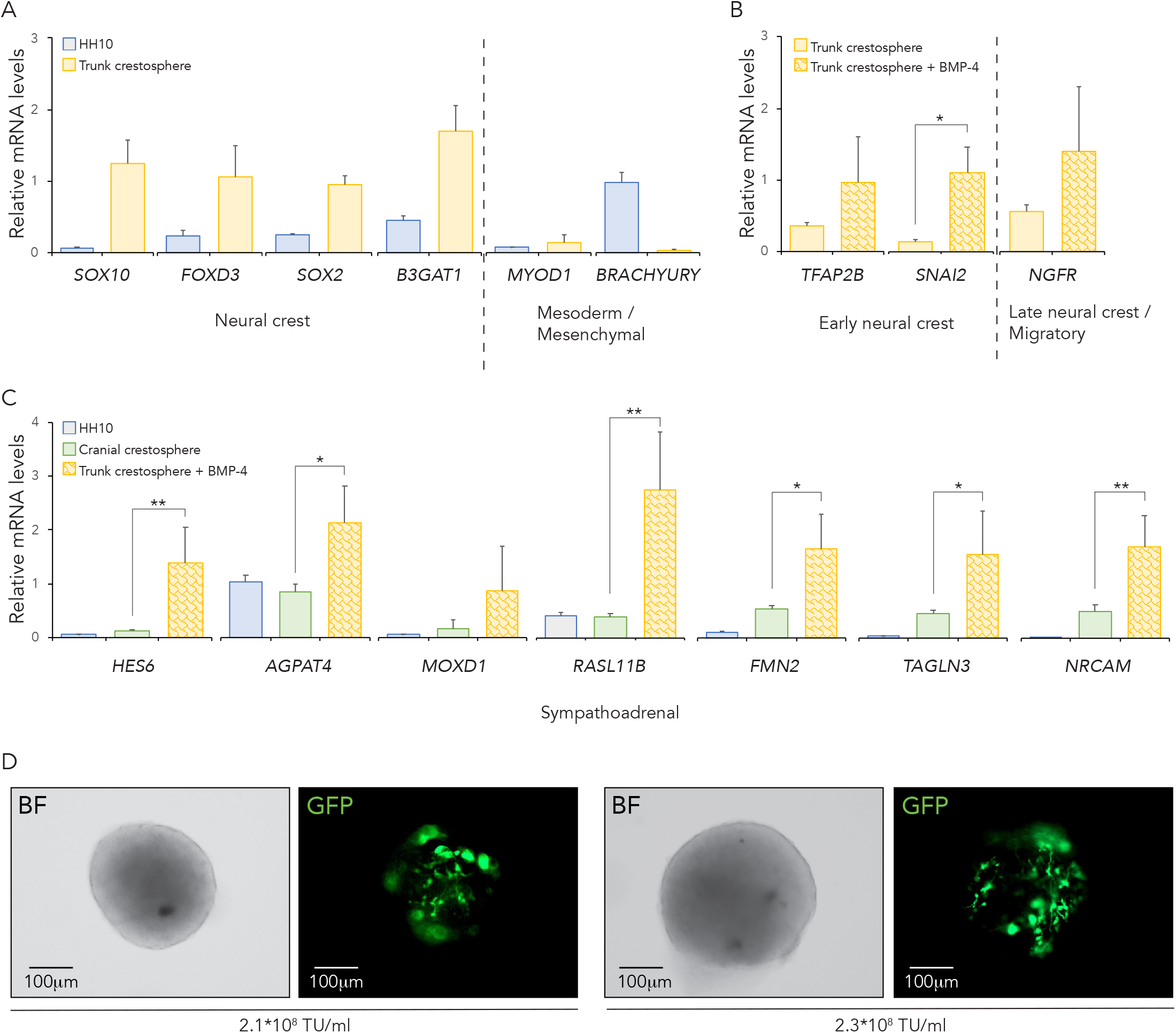
Quantitative PCR shows enriched neural crest gene expression in trunk crestospheres. **(A)** Relative mRNA expression of neural crest- and mesodermal/mesenchymal-associated genes comparing trunk crestosphere cultures (n=3) to stage HH10 wild type embryo (n=1, pool of three embryos). **(B)** Relative mRNA expression of neural crest-associated genes comparing trunk crestospheres cultured with (n=3) or without (n=3) BMP-4. **(C)** Relative mRNA expression levels of sympathoadrenal genes comparing trunk crestospheres cultured with BMP-4 (n=3) to HH10 wild type embryo (n=1, pool of three embryos) and cranial crestosphere cultures (n=4). **(A-C)** Data is presented as mean of n ± SD, and significance between trunk crestospheres cultured with or without BMP-4 (B) and trunk crestosphere and cranial crestosphere cultures (C) was calculated using two-sided student’s t-test; *p<0.05, **p<0.01, ***p<0.001. **(D)** GFP expression in individual trunk crestosphere cells transduced with different concentration of the virus shows similar transfection efficiency at all conditions. TU; Transducing Units.

*Addition of BMP-4 to the culture medium promotes neural crest maintenance and expansion* During normal development BMP-4 is secreted by the dorsal neural tube at the premigratory stage to promote neural crest fate (Liem et al., 1995), and later by the dorsal aorta, providing cues for neural crest cell migration as well as sympathoadrenal and chromaffin cell fate (Unsicker et al., 2013). We therefore compared expression of neural crest genes in trunk crestosphere cultures without (i.e. ‘*normal*’ NC medium) or with BMP-4 supplement, testing whether addition of BMP-4 could increase relative expression levels of neural crest genes. By visual inspection, the crestospheres grew similarly with or without BMP4 (Figure 2A). However, expression of the early neural crest marker genes *TFAP2B* and *SNAI2* increased (Figure 3B), and we also detected increased expression of *NGFR*, a marker expressed later by migrating and maturing peripheral neurons (Heuer et al., 1990) (Figure 3B). We conclude that BMP-4 further enriched neural crest fate in the cultures and should be routinely added to the optimized trunk crestosphere culture medium (Table 1). Altogether, we show that increased concentrations of RA are crucial for trunk crestosphere culture expansion (Table 1), and that BMP-4 enhances expression of at least some neural crest marker genes (Figure 3B).

### Trunk-derived crestospheres consist of a combination of neural crest stem and progenitor cells

Next, we tested whether the trunk crestospheres consisted of a pure stem cell population of premigratory neural crest cells, or as shown in many spheroid cultures of different stem cell types, a mixture of cells at multiple developmental stages (Jensen and Parmar, 2006; Piscitelli et al., 2015). To this end, we performed qPCR to determine relative expression of genes associated with more progenitor-like neural crest cells including markers of the migratory stage as well as genes reflecting the sympathoadrenal lineage. The results show that trunk crestospheres contain a mixture of neural crest stem-like cells as well as progenitor cells (Figure 3C). In addition, we observed enriched expression levels of recently characterized genes reflecting trunk neural crest, like *HES6* (Murko et al., unpublished), in trunk crestospheres as compared to cranial crestospheres, reflecting the posterior axial levels of the trunk crestospheres (Figure 3C).

### Trunk crestospheres can be efficiently transduced using lentiviral vectors

Genetic manipulation of crestospheres is one potential application for analysis on downstream effects of genes of interest (Dupin et al., 2018; Kerosuo and Bronner, 2016). However, our previous attempts using siRNAs have demonstrated low transfection efficiency (Kerosuo and Bronner, 2016). Therefore, we sought to determine whether the trunk-derived crestospheres could be efficiently transduced using viral vectors to create stable phenotypes. In a proof-of-principle experiment, we used a GFP-tagged lentiviral vector (pCMV-IRES-GFP version 3; pCIG3) and transduced trunk crestospheres with increasing concentrations of the virus. Trunk crestospheres did not display toxic side effects after viral transduction and survived well at all concentrations investigated. By analyzing GFP expression in individual spheres using live fluorescent microscopy, we observed no major differences in GFP intensity between the different viral concentrations tested (Figure 3D). These results show that trunk crestospheres can be efficiently transduced to create stable genotypes, overcoming the transient and inefficient nature of siRNAs.

### Conclusions

Crestospheres are an excellent tool to study different aspects of neural crest stemness and differentiation (Dupin et al., 2018; Kerosuo and Bronner, 2016; Kerosuo et al., 2015). Given the known axial level differences in the ability of cranial and trunk neural crest cells to give rise to different cell types (Ayer-Le Lievre and Le Douarin, 1982; Simoes-Costa and Bronner, 2016), here we optimized previous cranial crestosphere conditions (Kerosuo et al., 2015) for maintenance of trunk-derived neural crest cultures that arise from more posterior axial levels (Figure 4). Our protocol for trunk crestosphere cultures results in enrichment of neural crest specific genes as shown by *in situ* hybridization, immunostaining and qPCR, with little expression of mesodermal or mesenchymal markers.

Trunk crestospheres are not only a useful tool for studying normal neural crest development, but also promise to be useful for advancing research on neural crest-derived diseases and malignancies, including neuroblastoma and pheochromocytoma, as well as syndromes like Congenital Central Hypoventilation Syndrome (CCHS), Familial Dysautonomia (FD), or von Hippel-Lindau Syndrome ((Huber et al., 2018; Tsubota and Kadomatsu, 2018; Zhang et al., 2010). Given the ability to perform viral transduction, trunk crestospheres promise to provide insight into the molecular mechanisms that govern normal development as well as diseases derived from the more posterior parts of the embryo.

**Figure 4.**
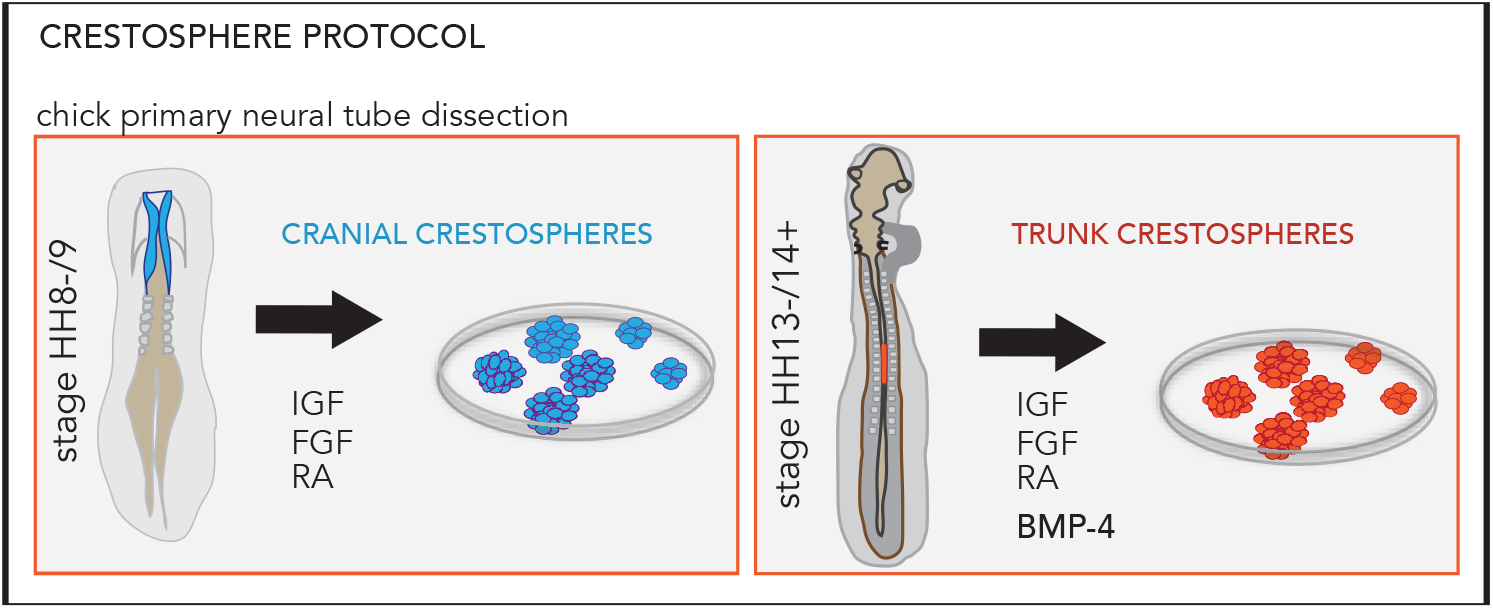
Schematic figure of respective axial levels dissected for cranial and trunk crestospheres, and the main differences in the culture medium composition.

## Acknowledgements

This work was supported by the Swedish Childhood Cancer Foundation (to SM), the Mary Bevé Foundation (to SM), Magnus Bergvall’s Foundation (to SM), the Thelma Zoéga Foundation (to SM), Hans von Kantzow Foundation (to SM), the Royal Physiographic Society of Lund (to SM), NIH Ruth L. Kirschstein NRSA F32HD087026 (to EK), NIH R01DE024157 (to MB), the Academy of Finland (to LK), Sigrid Juselius Foundation (to LK), and in part by the Division of Intramural Research of the National Institute of Dental and Craniofacial Research at the National Institutes of Health, Department of Health and Human Services.

